# Poly(A) tail length regulation by mRNA deadenylases is critical for suppression of transposable elements

**DOI:** 10.1101/2023.03.23.533991

**Authors:** Ling Wang, Hui Li, Zhen Lei, Mengxiao Yan, Yuqin Wang, Jiamin Zhao, Hongxia Wang, Jun Yang, Jungnam Cho

## Abstract

Transposons are mobile genetic elements that can impair the host genome stability and integrity. In plants, suppression of transposons is thought to be mediated mainly by small RNAs; however, the role of RNA decay in posttranscriptional repression of transposons is unknown. Here we show that RNA deadenylation is critical for controlling transposons in *Arabidopsis*. Previously, we demonstrated that transposon RNAs often harbor structural aberrancy owing to its inherently suboptimal codon usage and ribosome stalling. Such RNA aberrancy is monitored and resolved by RNA decay which is initiated by removal of poly(A) tail or deadenylation. The CCR4-NOT complex is a primary RNA deadenylase in *Arabidopsis*, and we found that it is required for stable repression of transposons. Intriguingly, RNA deadenylation controls transposons that are not targeted by cytoplasmic secondary small RNAs, which implies a target-specific regulation of transposon by the host. Our study suggests a previously unknown mechanism for transposon repression mediated by RNA deadenylation and unveils a complex nature of the host’s strategy to maintain the genome integrity.

## Introduction

Transposable elements (TEs) are mobile genetic elements that pose a significant threat to the host genome stability and integrity. It is well documented that transposons are subject to epigenetic silencing that is mediated by so-called RNA-directed DNA methylation (RdDM)^1^. Transposons that escape such transcriptional suppression or are newly introduced to the host genome, thus are yet epigenetically silenced, are recognized by the RNA-DEPENDENT RNA POLYMERASE 6 (RDR6)-SUPPRESSOR OF GENE SILENCING 3 (SGS3) complex which generates 21/22-nucleotide (nt) small interfering (si) RNAs to initiate epigenetic silencing^2–5^. Accumulating evidence suggests that the incompleteness of mRNA (e.g. removal of poly(A) tail or RNA truncation) is crucial for specific targeting of RDR6^2,6,7^. In our previous study, we showed that the suboptimal codon usage of transposons cause ribosome stalling and RNA cleavage, which accounts for their frequent targeting to the RDR6-mediated siRNA biogenesis pathway^8^. In addition, the ribosome-stalled transcripts are preferably guided to cytoplasmic RNA granules (RGs), for which the liquid-liquid phase separation of SGS3 is critical^8–10^. It is, however, important to note that the 21/22-nt siRNAs are associated with only around one third of active and transcribed TEs in *Arabidopsis*, and the posttranscriptional suppression of transposon that is independent of siRNAs is largely unknown.

Aberrancy of mRNA such as premature termination and arrest of translation is resolved by the RNA decay pathways^11–15^. mRNA deadenylation is a primary and rate-limiting step of RNA decay and is catalyzed by multiple deadenylase complexes^16^: the POLY(A) NUCLEASE 2 (PAN2)-PAN3 complex acts at an early phase of deadenylation in metazoa and yeast, and degrades the poly(A) tail to 50-110 nt^16–21^. However, orthologues of PAN2-PAN3 have not been identified in flowering plants^22,23^. The CARBON CATABOLITE REPRESSION 4 (CCR4)-NEGATIVE ON TATA-LESS (NOT) complex catalyzes more rapid deadenylation by two catalytic components, CCR4 and CCR4-ASSOCIATED FACTOR 1 (CAF1)^16,17^. In *Arabidopsis*, CCR4a, CCR4b, CAF1a, and CAF1b show catalytic activity of deadenylation^24–27^. A third class of deadenylase is POLY(A)-SPECIFIC RIBONUCLEASE (PARN), the orthologues of which have been identified in vertebrates and plants^16,23^. Loss of *PARN* genes in *Arabidopsis* causes embryo lethality^28,29^.

Despite the inherent aberrancy of transposon transcripts, their regulation by the cellular RNA surveillance system has been scantily investigated. A rather indirect evidence was provided in a recent study suggesting that the *Drosophila* mutant defective in *CCR4* accumulates TE transcripts, and the CCR4-NOT complex interacts with the piRNA pathway components in the nucleus, indicating a co-transcriptional suppression of transposon by a nuclear RNA deadenylation factor^30^. However, it has not been fully elucidated if the nuclear function of CCR4 in *Drosophila* requires the RNA deadenylation activity and the poly(A) tail regulation in transposon repression has never been studied in plants. In this study, we investigated the mutants for RNA deadenylases in *Arabidopsis* and assessed the transposon RNA levels. Intriguingly, we found that RNA deadenylases suppress a set of transposons that are not usually regulated by the RDR6-mediated pathway. Oxford Nanopore direct RNA sequencing (ONT-DRS) revealed that CCR4a shortens the poly(A) tails, destabilizes the transcripts, and reduces the steady-state mRNA levels of transposons. Moreover, we also carried out whole-genome resequencing and droplet digital PCR (ddPCR) experiments to interrogate the mobilization of TEs and observed a marked increase of transposon mobility in the deadenylase mutants. Our study unveils a previously unknown cellular mechanism that degrades transposon RNAs through an evolutionarily conserved RNA surveillance system.

## Results

### mRNA deadenylases suppress transposon expression

We previously showed that *Arabidopsis* TE RNAs often undergo ribosome stalling and RNA cleavage^8^. Since such aberrancy of RNA is monitored and resolved by RNA surveillance and decay pathways^15,17,31–33^, we reasoned that transposon RNAs might be controlled by the RNA degradation pathways. To test this possibility, we first identified the *Arabidopsis* mutants for RNA deadenylases (*ccr4a-1* and *ccr4b-1* single mutants, and *caf1a-1 caf1b-3* double mutant) and induced *de novo* mutations in *DECREASE IN DNA METHYLATION 1* (*DDM1*) using CRISPR-Cas9 to release transposons from epigenetic silencing (Supplementary Figure 1). It is worth noting that the pre-existing *ddm1* mutants contain significant number of newly inserted transposons that could be unevenly segregated in genetic crosses with other mutants, and therefore may lead to erroneous assessment of transposon expression. For this reason, we generated *de novo* mutants of *DDM1* and used the plant materials collected in the same generation (See Materials and methods). RNA-seq was then carried out in two independent *ddm1* mutant alleles of each RNA deadenylase mutant (Supplementary Figure 2). The transcriptome data showed a comparable number of genes that are up- and down-regulated; however, transposons exhibited a strikingly different pattern that most differentially expressed transposons are up-regulated in the deadenylase mutants (Fig. 1a-c). We then compared the up-regulated transposons in these three mutants and found that a large fraction of TEs is commonly up-regulated, while *CCR4a* displays the greatest impact on transposon expression (Fig. 1d and e; Supplementary Figure 3). These data imply that the mRNA deadenylation pathway is involved in transposon repression.

**Fig. 1.**
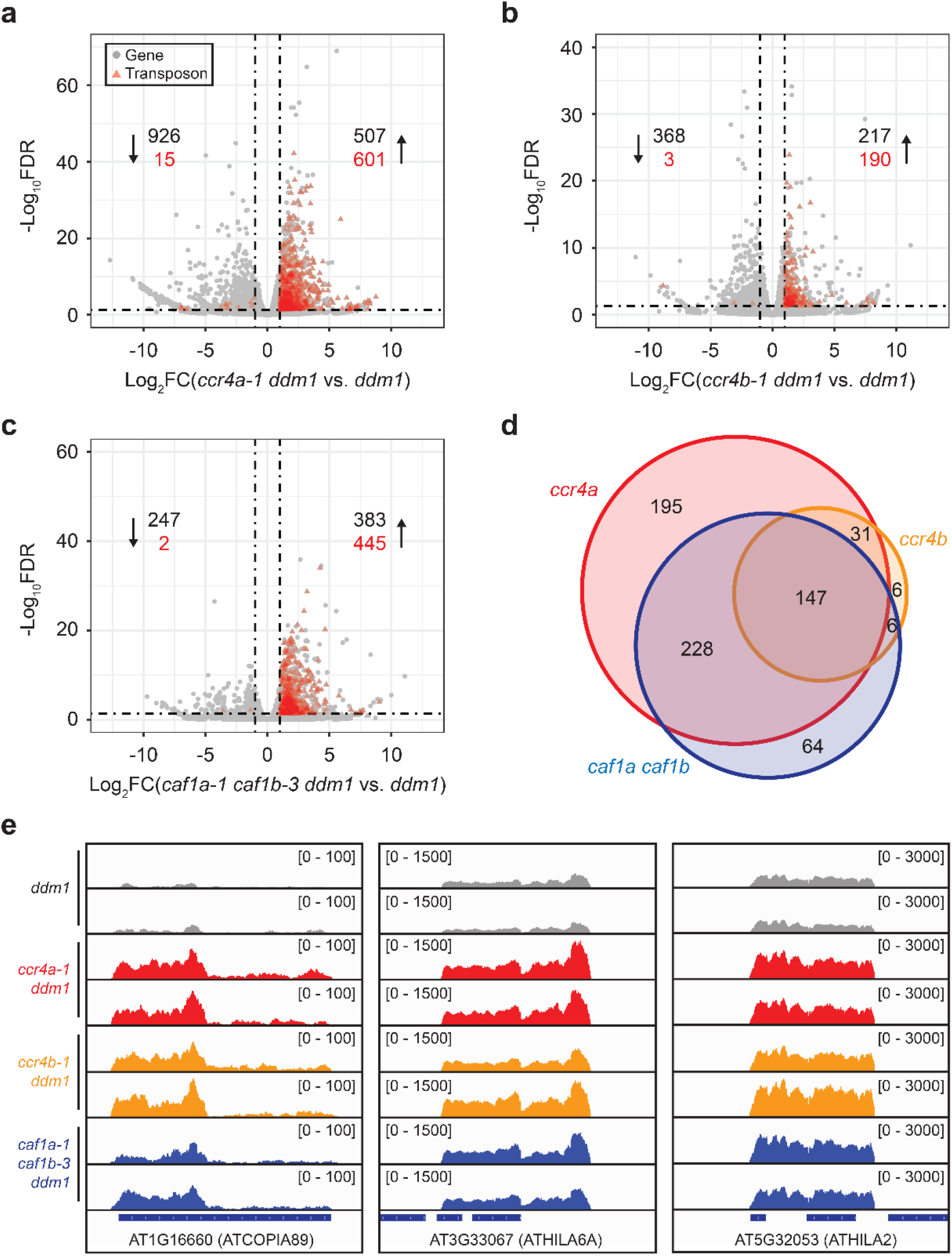
Loss of mRNA deadenylases leads to transposon derepression. **a-c** Volcano plots shown for *ccr4a-1 ddm1* (**a**), *ccr4b-1 ddm1* (**b**), and *caf1a-1 caf1b-3 ddm1* (**c**) in comparison with the *ddm1* single mutant. Mutations for *DDM1* was generated *de novo* by CRISPR-Cas9, and two independent *ddm1* mutant lines were used. Differential expression was defined by the log2-fold change greater than 1 or less than -1 and FDR values less than 0.05. Grey dots and red triangles represent genes and transposons, respectively. Numbers indicate differentially expressed genes and transposons, and up- or down-regulation was expressed by arrows. **d** Overlap of transposons upregulated by the mutations of *CCR4a, CCR4b*, and both *CAF1a* and *CAF1b*. **e** Genome browser snapshots for representative transposon loci showing the increased expression levels in the mRNA deadenylase mutants. Numbers in parentheses indicate read coverage and two independent *ddm1* mutant lines are displayed in separate tracks.

### Divergence of TE regulation by RDR6 and CCR4a

It is well documented that active transposons give rise to 21/22-nt siRNAs that can target transposon RNAs for cleavage^2,5^. Since the cleaved transcript products are eliminated by the cellular RNA decay pathways, we suspected that the increased expression of transposons in the deadenylase mutants might be due to compromised RNA decay of the cleaved transcripts by the RDR6-generated secondary siRNAs. However, the transposons regulated by CCR4a only marginally overlapped with those targeted by RDR6 (Fig. 2a), suggesting that the RNA deadenylases control different set of transposons and act in a siRNA-independent manner. Transposon classification analysis further supports this conclusion; the *RDR6*-regulated transposons are strongly enriched with the LTR/Gypsy family, and in the *ccr4a* mutants, DNA/MuDR DNA transposon family is strongly overrepresented (Fig. 2b). To further confirm the divergence of the RDR6- and CCR4a-regulated transposons, we compared the 21/22- and 24-nt siRNA levels. As shown in Fig. 2c, the 21/22-nt siRNAs of RDR6-controlled TEs were greatly increased in *ddm1*, whereas CCR4a-regulated transposons exhibited significant reduction of both classes of siRNAs in *ddm1*. In addition, the transposons regulated by RDR6 and CCR4a were mapped across the *Arabidopsis* chromosomes. The RDR6 target transposons were mostly found in the centromeric region, and the transposons regulated by CCR4a were also mapped to the pericentromeric and euchromatic regions in addition to centromeres (Fig. 2d; Supplementary Figure 4). Collectively, these data indicate that the mRNA deadenylases suppress transposon expression in a siRNA- and RDR6-independent manner.

**Fig. 2.**
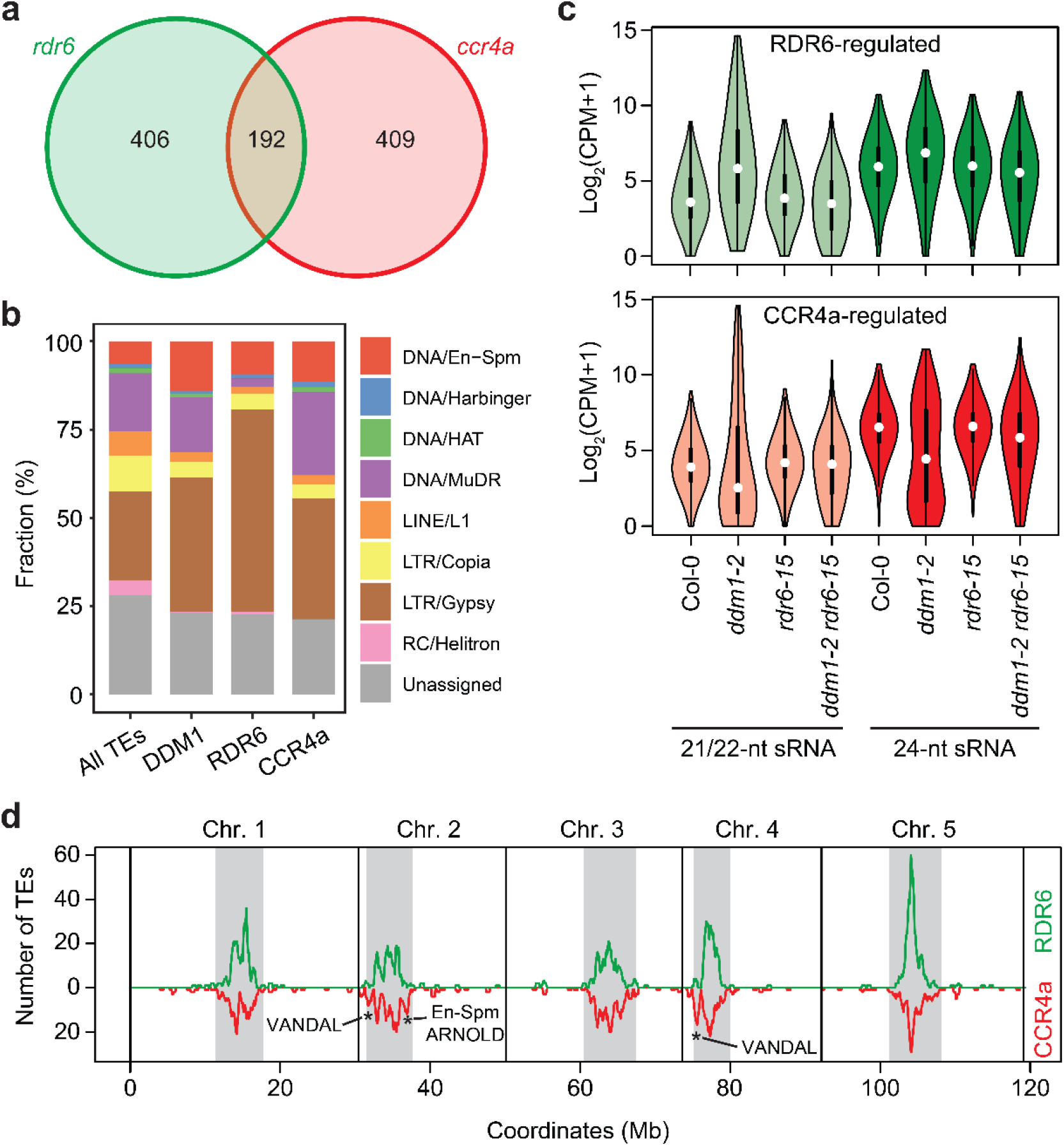
CCR4a regulates distinct set of transposons from those controlled by RDR6. **a** Overlap of transposons regulated by RDR6 and CCR4a. RDR6-regulated transposons were retrieved from the previous study^8^ and identified by the reduced 21/22-nt siRNA levels in the *rdr6 ddm1* double mutant as compared with the *ddm1* single mutant. CCR4a-regulated transposons are those identified in Fig. 1a. **b** Fraction of transposon families in all annotated transposons, derepressed in *ddm1*, regulated by RDR6, and upregulated in the *ccr4a* mutant. Reactivated transposons in the *ddm1* mutant were identified from a public dataset (GSE52952) by filtering those with the log2-fold change greater than 1 and FDR lower than 0.05. **c** Levels of 21/22- and 24-nt sRNAs in the transposons regulated by RDR6 and CCR4a. White circle, median; black rectangle, upper and lower quartile. The sequencing datasets were obtained from GSE52952. **d** Chromosomal distribution of RDR6- and CCR4a-regulated transposons. Numbers of TEs in 500-kb overlapping windows sliding in steps of 100 kb are shown. Regions overrepresented in the CCR4a-regulated transposons are marked by asterisks, and representative transposon families corresponding to the region are also indicated. Pericentromeric regions are expressed as grey boxes.

### CCR4a shortens poly(A) length and destabilizes transposon RNAs

We next wanted to measure the poly(A) tail lengths of TE RNAs in the deadenylase mutant. For this, we took advantage of ONT-DRS which allows for direct tail length measurement of native RNA. Transposon expression determined by ONT-DRS reproducibly showed a strongly increased expression in the *ccr4a* mutant (Supplementary Figure 5), verifying our observation shown in Fig. 1. The ONT-DRS data from *ddm1-L2* showed tail length peaks at 20, 40-50, and 70-80 nt, which are distanced by ∼25 nt (Fig. 3a and b). A similar pattern was also observed in previous studies^34,35^, suggesting a robust estimation of poly(A) tail length by ONT-DRS. Importantly, the *ccr4a-1 ddm1-L2* mutant displayed a considerably longer tail length compared to *ddm1-L2*, confirming that CCR4a is a key factor shortening the poly(A) tail. We then retrieved the transposon transcripts from our ONT-DRS dataset and analyzed their tail lengths. As shown in Fig. 3c and d, TE RNAs possess longer poly(A) tails compared to non-TE transcripts, and the *ccr4a* mutation resulted in by far longer tails.

**Fig. 3.**
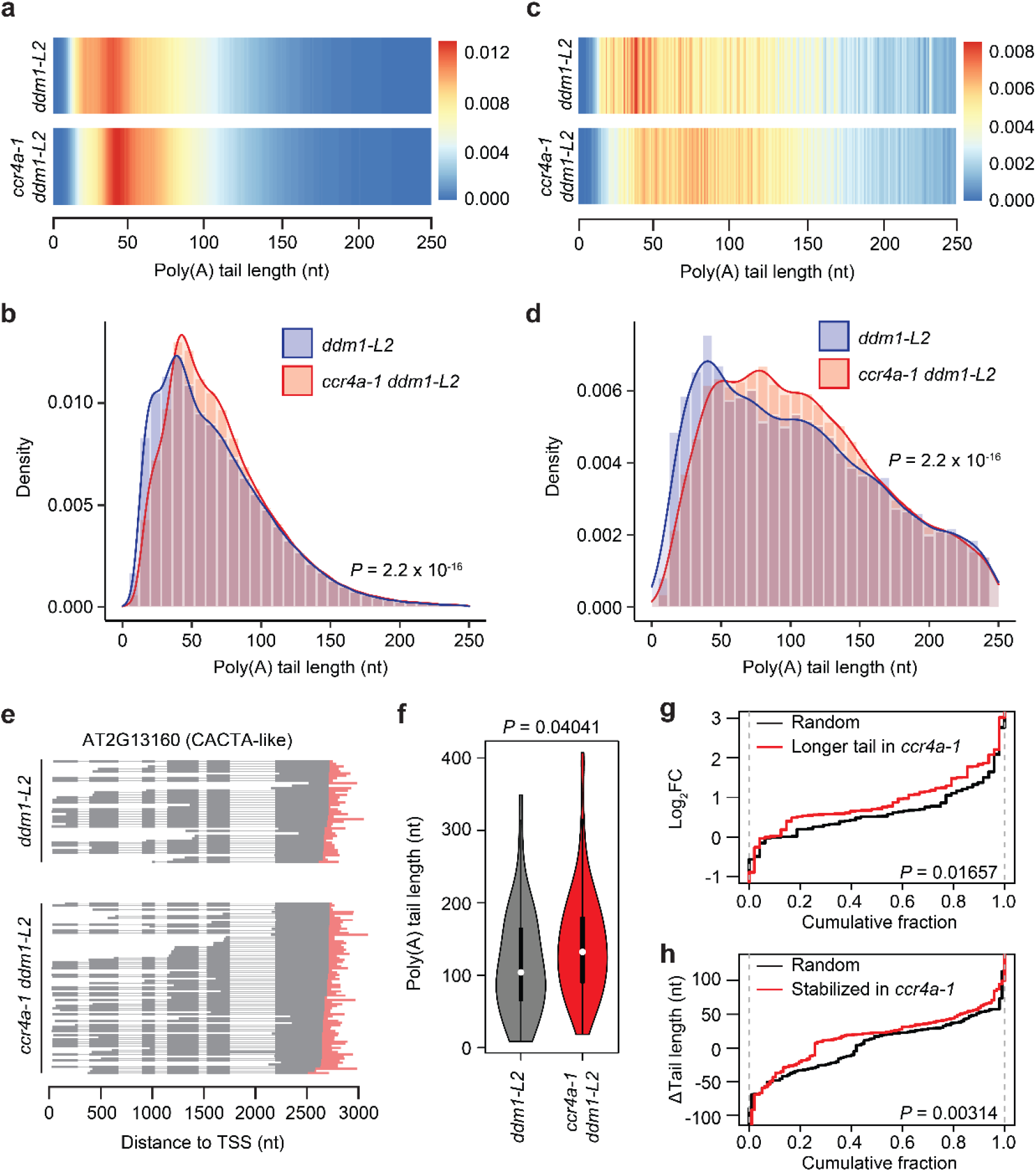
Longer transposon RNA tails are associated with increased expression. **a-d** mRNA tail lengths of all transcripts (**a-b**) and transposon RNAs (**c-d**) in *ddm1-L2* and *ccr4a-1 ddm1-L2*, shown as heatmap (**a** and **c**) and density plot (**b** and **d**). Poly(A) length was measured by Oxford Nanopore direct RNA sequencing. *P* value was obtained by the one-sided Wilcoxon rank sum test. **e-f** A CACTA-like TE exhibiting higher expression (**e**) and longer tail length (**f**) in *ccr4a-1 ddm1-L2* double mutant. In **e**, each line represents individual transcript detected by Oxford Nanopore direct RNA sequencing, and poly(A) tail is shown in red line. In **f**, mRNA tail lengths of the CACTA-like element in *ddm1-L2* and *ccr4a-1 ddm1-L2* are compared. White circle, median; black rectangle, upper and lower quartile. *P* value was obtained by the one-sided Wilcoxon rank sum test. **g** Fold changes of CCR4a-regulated transposons (n=48, log2-fold change of *ddm1-L2* vs. *ccr4a-1 ddm1-L2*) compared with randomly chosen transposons (n=48). CCR4a-regulated transposons are those with longer tails by at least 10 nt in the *ccr4a-1 ddm1-L2* mutant. *P* value was obtained by the one-sided Wilcoxon rank sum test. **h** Poly(a) tail length difference of transcripts stabilized by the loss of *CCR4a* in *ddm1-L2* and *ccr4a-1 ddm1-L2*. mRNA half-lives were determined for *ddm1-L2* and *ccr4a-1 ddm1-L2* by the transcription arrest assay followed by RNA-seq. Transcripts with longer half-lives in *ccr4a-1 ddm1-L2* by at least 0.5 h were selected (n=97) and compared against randomly chosen transcripts (n=100). Tail length difference was calculated by subtracting the tail lengths in *ddm1-L2* from those in *ccr4a-1 ddm1-L2. P* value was obtained by the one-sided Wilcoxon rank sum test.

It has been previously reported that highly expressed and stable mRNAs are featured with short steady-state poly(A) tail length^16,34^. Nonetheless, lengthening poly(A) tail contributes to active translation and RNA stability in humans and plants^25,36^. For example, the poly(A) tail length of a CACTA-like transposon became longer, and its expression level was increased in the *ccr4a-1 ddm1-L2* mutant compared to *ddm1-L2* (Fig. 3e and f). We further tested other TE transcripts that have longer tails in the *ccr4a-1 ddm1-L2* mutant for their expression levels and found that almost 90% of these TEs gained expression levels in *ccr4a-1 ddm1-L2* (Fig. 3g). Moreover, a transcription arrest RNA-seq was carried out to determine the RNA stability in *ddm1-L2* and *ccr4a-1 ddm1-L2*. Genes that became more stabilized in *ccr4a-1 ddm1-L2* as compared with *ddm1-L2* (increased half-lives by at least 0.5 h) exhibited longer tail length when *CCR4a* is mutated (Fig. 3h). These data together suggest that CCR4a shortens the poly(A) tail and destabilizes transposon RNAs.

### CCR4a suppresses transposon mobilization

We have so far demonstrated that loss of RNA deadenylases results in the increased expression of transposons. This led us to test if transposons mobilize more strongly in the deadenylase mutants. To test this idea, we carried out a whole-genome resequencing experiment to interrogate transposon proliferation. Ten individual plants from each genotype (*ddm1, ccr4a-1 ddm1, ccr4b-1 ddm1*, and *caf1a-1 caf1b-3 ddm1*) were randomly chosen and analyzed for non-reference and neo-insertions of transposons using the SPLITREADER pipeline^37^. Intriguingly, all three RNA deadenylase mutants showed increased number of transposon insertions compared to the *ddm1* single mutant (Fig. 4a). We also found that the transposons that exhibited active mobilization are of largely different types from those transcribed in the mutants tested; for instance, the LTR/Gypsy type was among the most actively transcribed in *ddm1* and *ccr4a-1 ddm1* (Fig. 2b), but it was the LTR/Copia family that was most proliferative in these mutants (Fig. 4b). This indicates that additional layer of posttranscriptional regulation exists to control transposon mobilization. Moreover, a ddPCR experiment was carried out to validate the enhanced mobilization of transposons in the RNA deadenylase mutants. ddPCR is an experimental method that can quantitatively measure DNA copies and is particularly useful for assessing transposon copy number^38,39^. For this experiment, a representative LTR/Copia element in the *Arabidopsis* genome known as *Evade* was chosen because it was well characterized for its active mobilization in the epigenetic mutants. As shown in Fig. 4c, the copy number of *Evade* was greatly increased in *ccr4a-1 ddm1* compared to *ddm1* in both T4 and T5 generations, further supporting our conclusion that RNA deadenylation represses transposition. In short, the RNA deadenylases suppress transposon expression and thereby inhibit the subsequent step of mobilization in *Arabidopsis*.

**Fig. 4.**
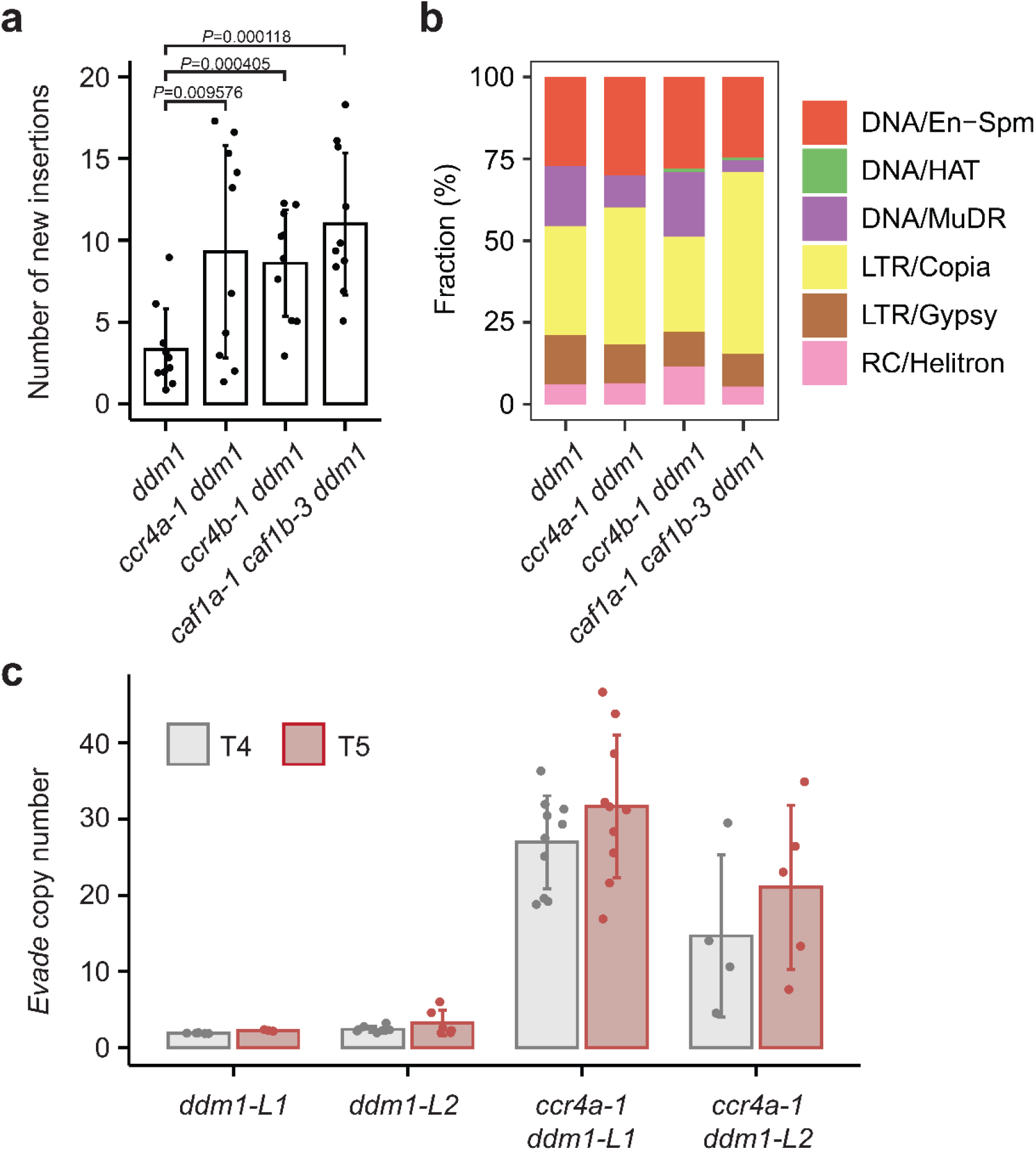
CCR4a suppresses transposon mobilization. **a** Number of new insertions of TEs detected in *ddm1, ccr4a-1 ddm1, ccr4b-1 ddm1*, and *caf1a-1 caf1b-3 ddm1*. 10 individual plants were randomly chosen, and whole-genome resequencing was performed for each individual plant independently. Data is presented in mean ± sd. *P* value was obtained by the one-sided Student’s t-test. **b** Percentage of TE families that were detected for neo-insertions. **c** Droplet digital PCR experiment determining the copy number of *Evade* retroelement in *ddm1* and *ccr4a-1 ddm1*. Plants were randomly chosen and extracted for DNA individually. The experiment was performed at T4 and T5 generations. Data is presented in mean ± sd.

## Discussion

In this study, we showed that RNA deadenylation is a critical mechanism that augments the host’s suppression of transposons in addition to the siRNA-mediated pathways. This suggests a previously unknown complexity of transposon control which is specific to TE types and sequences. We speculated that such divergence of transposon control might be attributed to differential subcellular localization of TE transcripts. Given that shortening of poly(A) tail is often coupled to translational repression^16,21^ and weak translation leads to localization to RNA granules which contain RNA deadenylases and degrading enzymes^8,22,24,40^, the TE transcripts controlled by RNA deadenylases might be more strongly enriched in RNA granules. Indeed, in the comparison between the RDR6- and CCR4a-regulated transposons, we were able to observe that the transcripts controlled by CCR4a exhibit relatively weaker translational activity and stronger RNA granule enrichment (Supplementary Figure 6).

Transposon repression by cytoplasmic siRNAs is considered an efficient, stable and perpetual way to control transposons because it can switch on the epigenetic silencing and target other TE transcripts with similar sequences^2,5^. On the other hand, RNA decay is merely degeneration of transcripts and does not generate any signals or molecules that can be amplified or transmitted. This partly accounts for why the RDR6-mediated pathway primarily acts on transposons, particularly those that are structurally intact and young^3^, while non-TE transcripts are predominantly controlled by RNA decay^41–44^. In this regard, the CCR4a-targeted TEs might be older in age compared to those regulated by RDR6, and have likely undergone more evolutionary sequence degeneration, which makes them less harmful to the host genome and thus less demanded for siRNA production. This suggests that the host genome employs diverse cellular mechanisms to control transposons at varying evolutionary ages.

Our work mainly focused on the cytoplasmic role of RNA deadenylases and directly demonstrated the poly(A) tail regulation of transposons using ONT-DRS. A previous report suggested a nuclear role of the CCR4-NOT complex in *Arabidopsis* as one of the essential elements for RdDM^45^. This is reminiscent of what is known in *Drosophila* that CCR4 co-transcriptionally represses transposon in association with Piwi^30^. These together suggest that the RNA deadenylase complex control transposons in the nucleus; however, it has not yet been elucidated if the nuclear function of the CCR4-NOT complex requires RNA deadenylation activity.

In summary, the shortening of transposon RNA poly(A) tail length by RNA deadenylases and thereby RNA destabilization is a critical cytoplasmic mechanism suppressing transposon activity. This work unveils a hidden complexity of transposon regulation, which helps broaden our understanding of the host’s defense against endogenous parasitic DNA.

## Materials and methods

### Plant materials and growth condition

All *Arabidopsis* plants used in this study are in the Col-0 background. The *ccr4a-1* (SAIL_802_A10), *ccr4b-1* (SALK_151541C), *caf1a-1* (SALK_070336), and *caf1b-3* (SALK_044043) mutants were obtained from the Arabidopsis Biological Resources Center (https://abrc.osu.edu/). The *caf1a-1 caf1b-3* double mutant was identified from the F2 segregation population derived from crosses. *De novo* mutants for *DDM1* were generated by CRISPR-Cas9 in the indicated mutant background, and the editing events were confirmed by Sanger sequencing. Unless otherwise stated, plants at T4 generation were used in this study. Sequences of primers used for genotyping are listed in Supplementary Table 1.

Seeds were sterilized using 75% ethanol, sown on Murashige and Skoog (MS) media [0.43 g/L MS salts (pH 5.8), 3 g/L sucrose, 0.8% (w/v) phytoagar] and stratified at 4 °C for 3 days. Plants were grown at 22 °C under long-day condition (16 h light/ 8 h dark)

### RT-qPCR

Total RNA was isolated from 10-day-old seedlings using the TRIzol extraction method (TIANGEN) and reverse-transcribed using ReverTra Ace qPCR RT Master Mix with gDNA Remover (TOYOBO). To quantify the relative abundance of transcripts, quantitative PCR was carried out using a ChamQ Universal SYBR qPCR Master Mix (VAZYME) on a CFX96 Touch Real-Time PCR Detection System (Bio-Rad). *Actin2* (AT3G18780) was used as an internal control for normalization. Gene expression levels were determined by the ΔΔCt method. Sequences of primers are listed in Supplementary Table 1.

### RNA-seq

For RNA-seq library construction, total RNA was isolated from 10-day-old seedlings using the Trizol Reagent (Invitrogen), and poly(A)-RNA was purified from 3 μg of total RNA using poly-T oligo-attached magnetic beads. Library was prepared using the NEBNext Ultra Directional RNA Library Prep Kit (NEB) following the manufacturer’s instructions. Sequencing was performed on an Illumina NovaSeq 6000 platform, and 150-bp paired-end (PE150) reads were generated. RNA-seq dataset is summarized in Supplementary Table 2.

For RNA-seq data analysis, the raw sequences were trimmed by Trimmomatic (version 0.39)^46^ to remove reads containing adapter and low-quality sequences with the parameters: LEADING:3 TRAILING:3 SLIDINGWINDOW:4:15 MINLEN:50.

Trimmed reads were then aligned to *Arabidopsis* reference genome (TAIR10) using Hisat2 (v.2.2.1)^47^ with settings: --rna-strandness RF --fr. Read count and FPKM values of genes and TEs were calculated by StringTie (v.2.1.7)^48^. The R package DESeq2^49^ was used for the differential expression analysis.

### Oxford Nanopore direct RNA sequencing (ONT-DRS)

Total RNA was isolated from 10-day-old seedlings by Trizol (Qiagen), and poly(A)-RNA was purified using Dynabeads mRNA Purification Kit (Invitrogen) following the manufacturer’s instructions. The quality and quantity of mRNA was assessed using the NanoDrop 2000 spectrophotometer and Qubit. Library was prepared using direct RNA sequencing kit (Nanopore, SQK-RNA002), loaded onto an R9.4 Flow Cell (Flow cell type FLO-MIN106), and sequenced on a GridION device for 72 hours. ONT-DRS dataset is summarized in Supplementary Table 3.

The raw nanopore signals were converted to base sequences by Guppy (v6.1.5) using the high-accuracy basecalling model. Since transposons are not properly annotated in the reference assembly and therefore often omitted in the downstream analysis, we merged all the de novo assembled transcripts derived from the RNA-seq data generated in this study and used the custom transcriptome assembly in our analysis.

Then, the nanopore reads with a mean quality score greater than 7 were mapped to the custom transcriptome using Minimap2 (v2.24-r1122)^50^ with the following parameters: -ax map-ont -L -p 0 -N 10. Poly(A) tail length was detected by Nanopolish (version 0.13.3)^51^. Transcripts with more than 15 reads were used to obtain the median poly(A) tail length. The reads with poly(A) tail were re-aligned to TAIR10 genome with the following parameters: -ax splice -k14 -uf and visualized by the python genome package bugv^34^.

### mRNA half-life

4-day-old etiolated *Arabidopsis* seedlings were immersed in cordycepin solution [1 mM PIPES (pH 6.25), 15 mM sucrose, 1 mM KCl, 1 mM sodium citrate, and 1 μM cordycepin] and harvested at 0, 0.25, 0.5, 1, 2, and 4 hrs for three biological replicates. RNA extraction and RNA-seq was performed as described above. mRNA half-lives were calculated as following: decay rate Ki = -ln(Fi/F0)/Ti, in which Fi is the FPKM at time i, and Ti is the time of cordycepin treatment. Ki was calculated from each time point, and the half-life is ln(2)/Ka, in which Ka is the average decay rate measured for all time points.

### Whole-genome resequencing

Genomic DNA was extracted using the CTAB method. 1 μg genomic DNA was randomly fragmented by ultrasonicator (Covaris). An average size of 200-400 bp DNA fragments were selected by Agencourt AMPure XP-Medium kit. The fragments were then end-repaired, 3’ adenylated and ligated with adaptors. The purified double-stranded products were heat denatured to single-stranded DNA, and then circularized. The single-stranded circular DNA was sequenced by a DNBSEQ-T7 generating 150 bp paired-end reads. Whole-genome resequencing dataset is summarized in Supplementary Table 4.

Paired-end short-read whole-genome sequencing data were mapped to TAIR10 and processed following the SPLITREADER pipeline^37^. Insertions supported by at least three reads (DP filter = 3) were filtered and only non-reference insertions were considered.

### Droplet digital PCR

Droplet digital PCR (ddPCR) was performed on TargetingOne® Digital PCR System (TargetingOne) following manufacturer’s instruction. Briefly, Genomic DNA was extracted using a N96 DNAsecure Plant Kit (Tiangen). 100 ng of genomic DNA was digested using AluI (NEB) for 4 h at 37 ° C. The digested DNA was quantified using the Qubit4 DNA quantification system (Thermo Fisher Scientific) and diluted to 0.15 ng/μL. The reaction mixture containing 2× ddPCR Supermix (Bio-Rad), 0.8 μM primer, 0.25 μM probe and 0.6 ng of cleaved sample DNA was thoroughly mixed and added into the droplet generation chip. Then, 180 μL of droplet generation oil was added to the mixture in the reaction mix inlet. Subsequently, the generated droplets were transferred into an 8-strip PCR tube and used for PCR reaction that was performed on a PTC-200 Thermal Cycler. FAM (488 nm) and VIC (532 nm) fluorescence signals were detected through the separate channels on the Chip Reader. Finally, the data were subjected to Poisson distribution analysis using the Chip Reader R1 software to obtain the target DNA copy numbers. Sequences of primers and probes are listed in Supplementary Table 1.

### Data availability

All the sequencing data generated in this study is available in the SRA repository under the accession ID PRJNA940263.

## Acknowledgements

This work was supported by the Strategic Priority Research Program of Chinese Academy of Sciences (XDB27030209), National Natural Science Foundation of China (32150610473, 32111540256, and 32270569) and General Program of Natural Science Foundation of Shanghai (22ZR1469100).

## Author contribution

JC conceived the idea and designed the experiments. HL, ZL, MY and YW conducted the experiments. LW, HL, HW, JY and JC analyzed the data. LW and JC drafted and wrote the manuscript. JC revised the manuscript.

