## Supplementary information for "Poly(A) tail length regulation by mRNA deadenylases is critical for suppression of transposable elements"

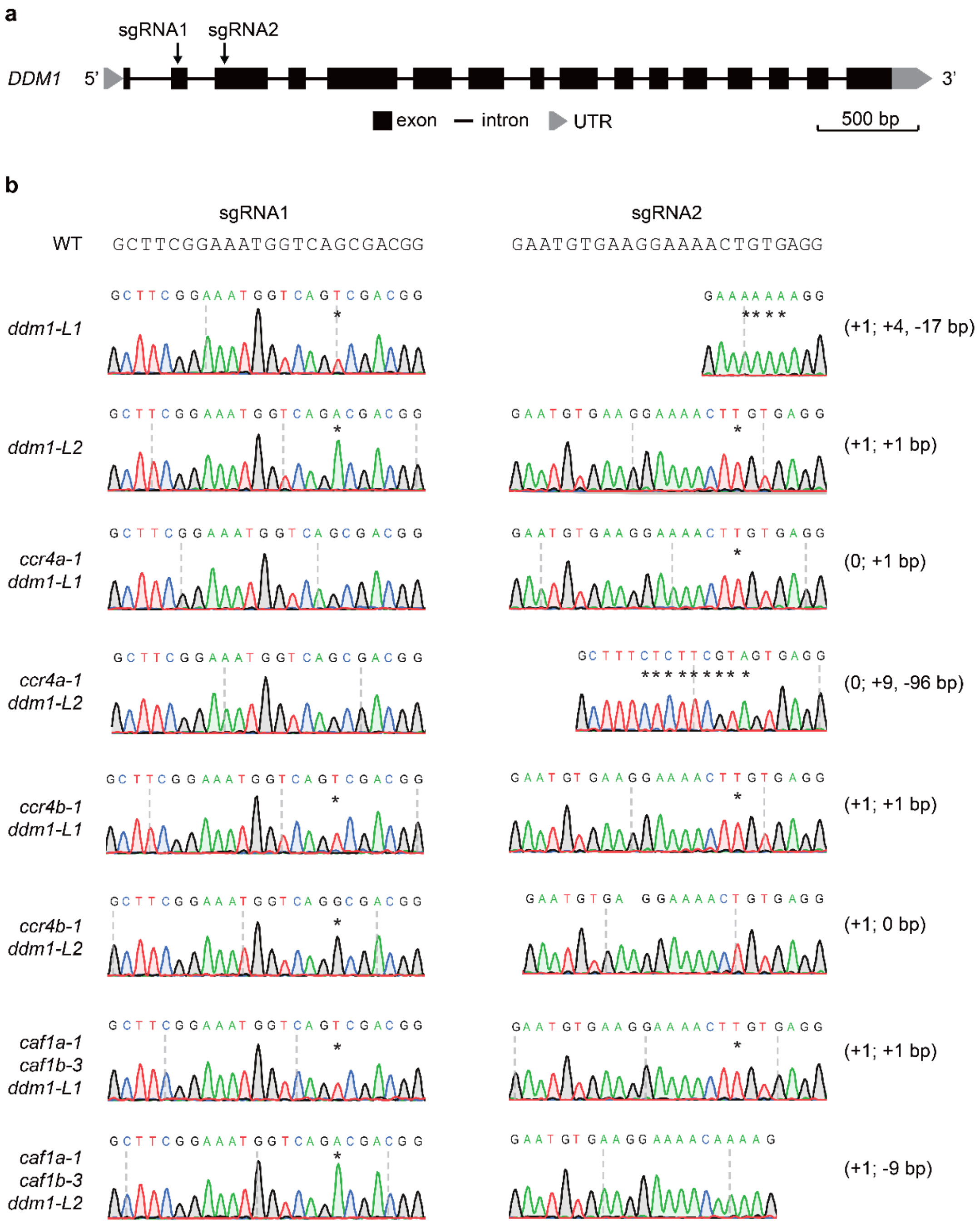


**Supplementary Fig. 1 | Identification of *de novo* mutated *DDM1* alleles.**

**a** Gene structure of *DDM1*. Two sgRNA-targeted sites are indicated.

**b** Chromatogram of Sanger sequencing around the targeted regions. Asterisks indicate the mutated sites. Numbers in parentheses denote nucleotide insertions and deletions.

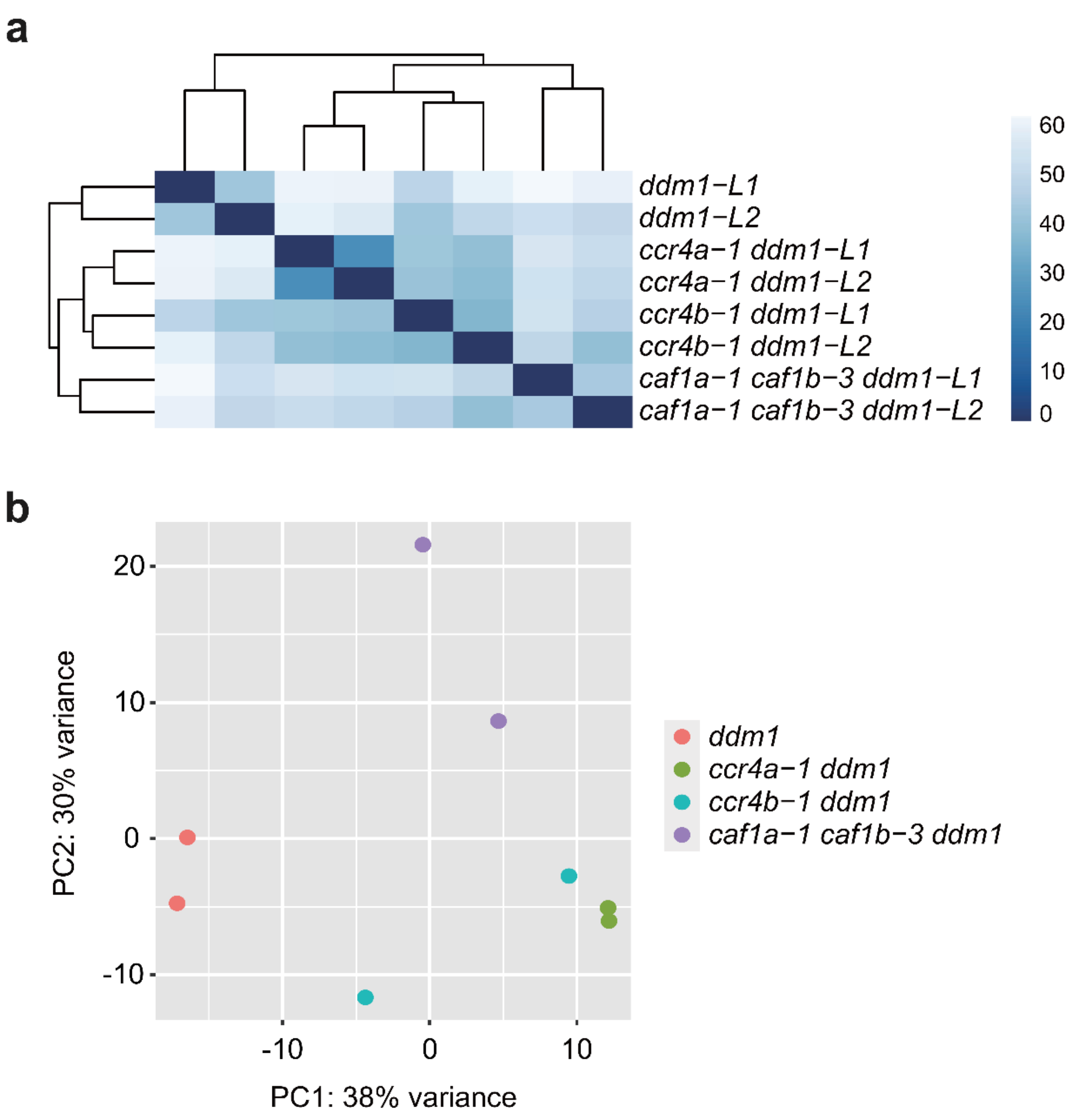


**Supplementary Fig. 2 | Similarity between two *ddm1* alleles in RNA-seq.**

**a** Distance matrix of *ddm1*, *ccr4a-1 ddm1*, *ccr4b-1 ddm1* and *caf1a-1 caf1b-3 ddm1*.

**b** Principal component analysis of *ddm1*, *ccr4a-1 ddm1*, *ccr4b-1 ddm1* and *caf1a-1 caf1b-3 ddm1*.

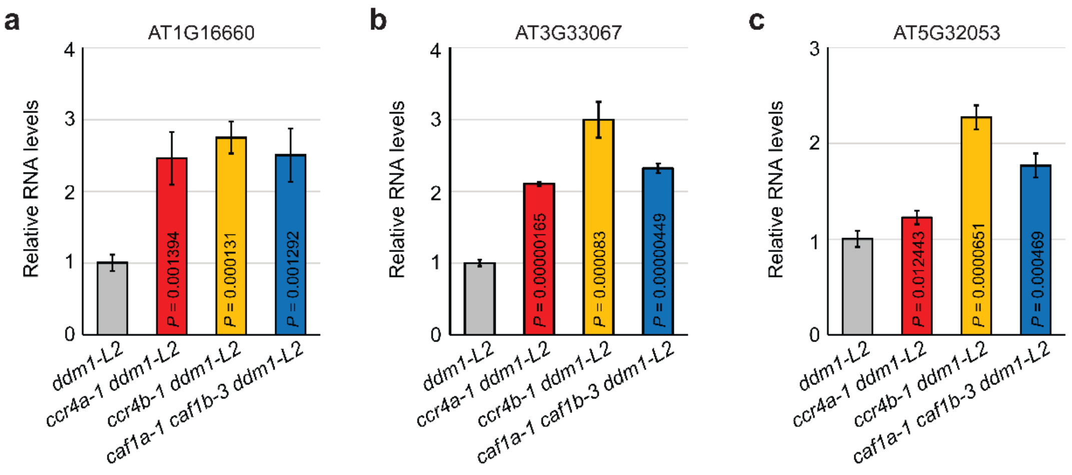


**Supplementary Fig. 3 | qPCR validation.**

**a-c** Relative expression levels of AT1G16660 (**a**), AT3G33067 (**b**), and AT5G32053 (**c**) determined by RT-qPCR. Data are shown in mean ± sd of three biological replications and *P* values are obtained in the comparison against *ddm1-L2* by the one-sided Student's t-test.

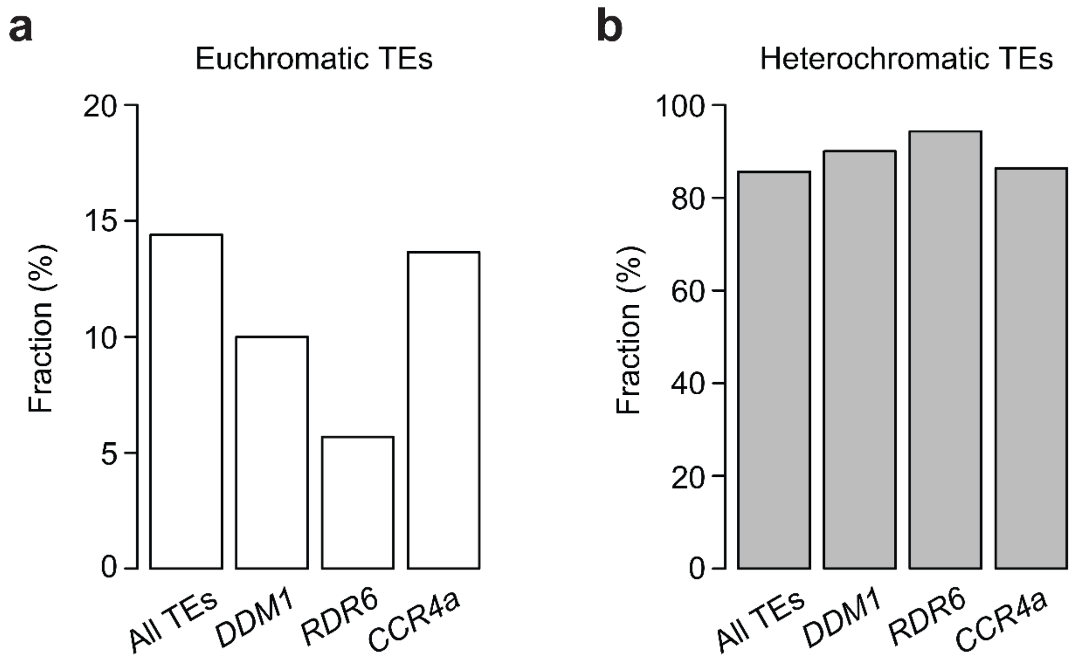


**Supplementary Fig. 4 | TE fractions in euchromatic and heterochromatin regions.**

**a** and **b** Fractions of TEs regulated by *DDM1*, *RDR6*, and *CCR4a* in the euchromatic (**a**) and heterochromatic (**b**) regions.

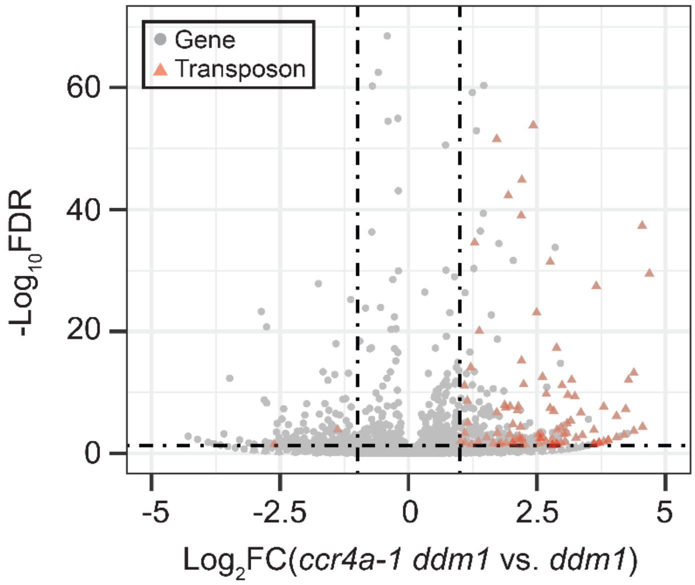


**Supplementary Fig. 5 | Reactivation of transposons in the *ccr4a* mutant.**

Volcano plot for *ddm1* and *ddm1 ccr4a-1* determined by Oxford Nanopore direct RNA sequencing.

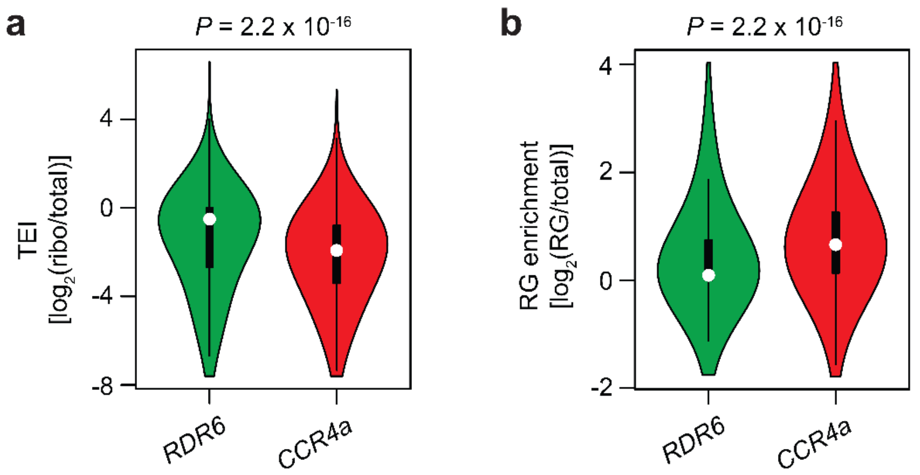


**Supplementary Fig. 6 | Translational efficiency and RNA granule enrichment.**

**a** and **b** Translational efficiency index (**a**) and RNA granule enrichment (**b**) of RDR6- and CCR4a-regulated transposons in *ddm1*. Transposons controlled by RDR6 and CCR4a were as defined in Fig. 2. TEI, translational efficiency index (log2-converted fold change of ribosome-protected to total RNA levels); RG enrichment, RNA granule enrichment (log2-convered fold change of RNA granule-enriched to total RNA levels). *P* values are obtained by the one-sided Wilcoxon rank sum test.

**Supplementary Table 1 | Sequences of primers used in this study.**

| Experiment | Primer name | Sequence (5' → 3') |
|---|---|---|
| **RT-qPCR** | Actin2-F1 | GGTAACATTGTGCTCAGTGGTGG |
| | Actin2-R1 | CAACGACCTTAATCTTCATGCTGC |
| | Evade-F1 | AATCGAAAGGGGGAGAAAGA |
| | Evade-R1 | CGCAAAACAAATGTCAGGTC |
| | AT1G16660-F | ATCAAGGCCACCAGCTTACC |
| | AT1G16660-R | GTAGGGGTCTGAGAGAGAGG |
| | AT3G33067-F | TCAGAGAGCACGTACCTCCA |
| | AT3G33067-R | GAGAAGCAGCTAAGCTTGTC |
| | AT5G32053-F | CCGCGGTTGCATAAACTACG |
| | AT5G32053-R | TAGAGAGCAGGAGATTCCGC |
| **ddPCR** | Evade-F2 | AGATTCTCCACGAAAGGCGT |
| | Evade-R2 | TAAGCAAGTGTTTAATTAGGTCATT |
| | Evade-FAM | FAM-ACCCTGGATTTAAGGTGAGAAGGAGTC-BHQ3 |
| | Actin2-F2 | GCTGATGATATTCAACCAATCG |
| | Actin2-R2 | CCTACCAACAACACTGGGAA |
| | Actin2-HEX | HEX-TGGTACCGGTATGGTGAAGGCTGG-BHQ3 |

**Supplementary Table 2 | Summary of RNA-seq.**

| Sample | Raw reads | Clean reads | Uniquely mapped reads | Alignment rate |
|---|---|---|---|---|
| *ddm1-L1* | 51052498 | 50585468 | 49082256 | 99.22% |
| *ddm1-L2* | 57040254 | 56545772 | 53470438 | 98.94% |
| *ccr4a-1 ddm1-L1* | 45709416 | 45418375 | 43853297 | 99.20% |
| *ccr4a-1 ddm1-L2* | 50680348 | 50337679 | 48648443 | 99.21% |
| *ccr4b-1 ddm1-L1* | 48335140 | 47887858 | 46342270 | 99.11% |
| *ccr4b-1 ddm1-L2* | 47842084 | 47464689 | 45873432 | 99.13% |
| *caf1a-1 caf1b-3 ddm1-L1* | 41499458 | 41131095 | 39298626 | 98.00% |
| *caf1a-1 caf1b-3 ddm1-L2* | 45544974 | 45164566 | 43662425 | 99.10% |
| *ddm1-L2* 0 h rep1 | 44831462 | 42953974 | 39103564 | 93.03% |
| *ddm1-L2* 0 h rep2 | 25951428 | 24872224 | 22757026 | 93.50% |
| *ddm1-L2* 0 h rep3 | 33762545 | 32279471 | 30130900 | 95.41% |
| *ddm1-L2* 0.25 h rep1 | 28129075 | 26850938 | 24518936 | 93.46% |
| *ddm1-L2* 0.25 h rep2 | 40336267 | 38627442 | 35279986 | 93.53% |
| *ddm1-L2* 0.25 h rep3 | 29632165 | 28549072 | 25973761 | 93.09% |
| *ddm1-L2* 0.5 h rep1 | 27635277 | 26597267 | 24270052 | 93.58% |
| *ddm1-L2* 0.5 h rep2 | 47071564 | 45049993 | 40991441 | 93.39% |
| *ddm1-L2* 0.5 h rep3 | 28076161 | 26919136 | 24573997 | 93.63% |
| *ddm1-L2* 1 h rep1 | 41268130 | 39108593 | 35642185 | 94.06% |
| *ddm1-L2* 1 h rep2 | 37307539 | 35611629 | 32231609 | 93.35% |
| *ddm1-L2* 1 h rep3 | 60329021 | 57548798 | 51904726 | 93.06% |
| *ddm1-L2* 2 h rep1 | 53305070 | 50668043 | 45467596 | 93.11% |
| *ddm1-L2* 2 h rep2 | 48349159 | 45976930 | 41360113 | 93.27% |
| *ddm1-L2* 2 h rep3 | 41732631 | 39667379 | 35670988 | 93.50% |
| *ddm1-L2* 4.25 h rep1 | 40198111 | 38192704 | 34274330 | 93.28% |
| *ddm1-L2* 4.25 h rep2 | 28280822 | 26845339 | 23915413 | 93.43% |
| *ddm1-L2* 4.25 h rep3 | 42331151 | 39858717 | 35922891 | 93.79% |
| *ccr4a-1 ddm1-L2* 0 h rep1 | 47988340 | 45721636 | 42164841 | 94.08% |

| | | | | |
|---|---|---|---|---|
| *ccr4a-1 ddm1-L2* 0 h rep2 | 27935915 | 26683639 | 24427322 | 93.37% |
| *ccr4a-1 ddm1-L2* 0 h rep3 | 45573618 | 43536708 | 39695594 | 93.04% |
| *ccr4a-1 ddm1-L2* 0.25 h rep1 | 38217233 | 36407324 | 33318949 | 93.54% |
| *ccr4a-1 ddm1-L2* 0.25 h rep2 | 35815137 | 33997702 | 31141120 | 93.82% |
| *ccr4a-1 ddm1-L2* 0.25 h rep3 | 28745163 | 27339148 | 22881859 | 93.52% |
| *ccr4a-1 ddm1-L2* 0.5 h rep1 | 38350730 | 36676662 | 33447055 | 93.42% |
| *ccr4a-1 ddm1-L2* 0.5 h rep2 | 34685255 | 33092235 | 30287348 | 93.78% |
| *ccr4a-1 ddm1-L2* 0.5 h rep3 | 40414492 | 38639145 | 35344818 | 93.73% |
| *ccr4a-1 ddm1-L2* 1 h rep1 | 53968763 | 51617566 | 47454617 | 94.50% |
| *ccr4a-1 ddm1-L2* 1 h rep2 | 45780313 | 43298155 | 39368821 | 93.46% |
| *ccr4a-1 ddm1-L2* 1 h rep3 | 37043983 | 35353693 | 32424847 | 94.18% |
| *ccr4a-1 ddm1-L2* 2 h rep1 | 37871391 | 36066939 | 32636252 | 93.43% |
| *ccr4a-1 ddm1-L2* 2 h rep2 | 51793971 | 49195941 | 44280490 | 93.83% |
| *ccr4a-1 ddm1-L2* 2 h rep3 | 56767076 | 54067756 | 48386973 | 93.02% |
| *ccr4a-1 ddm1-L2* 4.25 h rep1 | 42998405 | 40715120 | 36567534 | 93.12% |
| *ccr4a-1 ddm1-L2* 4.25 h rep2 | 39026849 | 36907240 | 33392188 | 93.55% |
| *ccr4a-1 ddm1-L2* 4.25 h rep3 | 46190008 | 43859830 | 39719118 | 93.65% |

**Supplementary Table 3 | Summary of Oxford Nanopore direct RNA sequencing.**

| Sample | Reads number (Q>7) | N50 | Aligned reads (primary) | Primary alignment rate |
|---|---|---|---|---|
| *ddm1-L2* | 1206306 | 998 | 1201102 | 99.57% |
| *ccr4a-1 ddm1-L2* | 1432587 | 1010 | 1423451 | 99.36% |

**Supplementary Table 4 | Summary of whole-genome resequencing.**

| Sample | Raw reads | Mapped reads | Alignment rate | Mean coverage |
|---|---|---|---|---|
| *ddm1* #1 | 26674266 | 25397090 | 95.21% | 31.8269 |
| *ddm1* #2 | 34309056 | 32487766 | 94.69% | 40.7126 |
| *ddm1* #3 | 29497002 | 28174863 | 95.52% | 35.3084 |
| *ddm1* #4 | 26635128 | 24987397 | 93.81% | 31.3123 |
| *ddm1* #5 | 25610746 | 24432721 | 95.40% | 30.6186 |
| *ddm1* #6 | 36347912 | 34722681 | 95.53% | 43.5128 |
| *ddm1* #7 | 29881272 | 28106678 | 94.06% | 35.2161 |
| *ddm1* #8 | 28480434 | 27227062 | 95.60% | 34.1193 |
| *ddm1* #9 | 30787622 | 29478403 | 95.75% | 36.9376 |
| *ddm1* #10 | 26564774 | 25427054 | 95.72% | 31.8636 |
| *ccr4a-1 ddm1* #1 | 24734532 | 23484789 | 94.95% | 29.4299 |
| *ccr4a-1 ddm1* #2 | 26662496 | 25512095 | 95.69% | 31.9696 |
| *ccr4a-1 ddm1* #3 | 31166140 | 29765882 | 95.51% | 37.302 |
| *ccr4a-1 ddm1* #4 | 26691168 | 25535428 | 95.67% | 31.9967 |
| *ccr4a-1 ddm1* #5 | 29894020 | 28653625 | 95.85% | 35.9102 |
| *ccr4a-1 ddm1* #6 | 26547840 | 25450769 | 95.87% | 31.8945 |
| *ccr4a-1 ddm1* #7 | 24667406 | 23640273 | 95.84% | 29.6256 |
| *ccr4a-1 ddm1* #8 | 27830820 | 26586169 | 95.53% | 33.3167 |
| *ccr4a-1 ddm1* #9 | 34125760 | 32249625 | 94.50% | 40.41 |
| *ccr4a-1 ddm1* #10 | 54113648 | 51639906 | 95.43% | 64.7133 |
| *ccr4b-1 ddm1* #1 | 37140154 | 35599292 | 95.85% | 44.6122 |
| *ccr4b-1 ddm1* #2 | 25552914 | 24562595 | 96.12% | 30.7799 |
| *ccr4b-1 ddm1* #3 | 24703268 | 23737461 | 96.09% | 29.7456 |
| *ccr4b-1 ddm1* #4 | 24714426 | 23723487 | 95.99% | 29.7298 |
| *ccr4b-1 ddm1* #5 | 25518522 | 24536020 | 96.15% | 30.7472 |

| | | | | |
|---|---|---|---|---|
| *ccr4b-1 ddm1* #6 | 28348814 | 27177916 | 95.87% | 34.0591 |
| *ccr4b-1 ddm1* #7 | 25590752 | 24343324 | 95.13% | 30.5039 |
| *ccr4b-1 ddm1* #8 | 33799022 | 32294517 | 95.55% | 40.473 |
| *ccr4b-1 ddm1* #9 | 29428316 | 28449248 | 96.67% | 35.6523 |
| *ccr4b-1 ddm1* #10 | 39053080 | 37477080 | 95.96% | 46.9689 |
| *caf1a-1 caf1b-3 ddm1* #1 | 30188638 | 26848607 | 88.94% | 33.6462 |
| *caf1a-1 caf1b-3 ddm1* #2 | 28171736 | 26932385 | 95.60% | 33.7478 |
| *caf1a-1 caf1b-3 ddm1* #3 | 24321696 | 23260323 | 95.64% | 29.1497 |
| *caf1a-1 caf1b-3 ddm1* #4 | 24927948 | 23696369 | 95.06% | 29.6964 |
| *caf1a-1 caf1b-3 ddm1* #5 | 29440560 | 27885587 | 94.72% | 34.9447 |
| *caf1a-1 caf1b-3 ddm1* #6 | 26859564 | 25227342 | 93.92% | 31.6126 |
| *caf1a-1 caf1b-3 ddm1* #7 | 27138548 | 25470911 | 93.86% | 31.9179 |
| *caf1a-1 caf1b-3 ddm1* #8 | 24688176 | 23306366 | 94.40% | 29.2051 |
| *caf1a-1 caf1b-3 ddm1* #9 | 26833208 | 25353700 | 94.49% | 31.773 |
| *caf1a-1 caf1b-3 ddm1* #10 | 38725746 | 36895528 | 95.27% | 46.2392 |
